# Prevalence of Meconium-Stained Amniotic Fluid and Its Associated Factors Among Postnatal Mothers In North Shewa Zone Public Hospitals, Oromia Ethiopia, 2023

**DOI:** 10.64898/2026.11.16.767185

**Authors:** Hawi Abayneh, Erean Shigign, Eleni Tesfaye, Mestawet Negash, Befikadu Tesfaye, Fekadu Tolosa, Gizu Tola, Misra Nurie, Jemal Bedane

## Abstract

**Background:** Meconium is a germ-free, dense, black-green, odor less substance that appears in the fetal intestine around 12 weeks of pregnancy and persists throughout the fetal colon during the pregnancy.. It is made up of 70% water, pancreatic and intestinal secretions, and desquamated epithelial cells from the intestinal tract and bile. In both developed and resource-limited countries, evidence showed that Meconium-stained amniotic fluid increased maternal and fetal morbidity and mortality. The prevalence and predictors of Meconium-stained amniotic fluid are not known in the North Shewa Zone. It is critical to understand the local burden and factors associated with MSAF for better intrapartum care and neonatal surveillance.

**Objective:** This study assessed prevalence of meconium-stained amniotic fluid and its associated factors among postnatal mothers in North Shewa Zone Public Hospitals, Oromia Ethiopia.

**Methods:** Institution based cross-sectional study was carried out among 495 postnatal mothers. Simple random sampling technique was used to select study participants. Pretested and structured questionnaires were used to collect data. Data were entered into Epidata and exported to SPSS for analysis. Bivariate and multi variable logistic regression was employed to identify an association between the independent predictors and the outcome variable.

**Results:** In this study, the proportion of Meconium stained amniotic fluid was found to be 29.1% (95%CI; 25.3-33.3). Inter-pregnancy interval, Gestational age, Prolonged premature rapture of membrane, Pregnancy induced hypertension, Duration of labor >15 hrs, Obstructed labor, Oligohydraminos and Current alcohol use were significantly associated with Meconium stained amniotic fluid.

**Conclusion:** Health education for the mothers during preconception care and antenatal care for the important early detection and disease prevention. Early detection by using a latent follow-up chart and partograph and timely intervention are recommended to decrease prolonged labor.

## Introduction

In Greek, Meconium refers to opium or poppy-like substances which found in the intestinal tract prior to birth. Meconium is a germ-free, dense, black-green, odor less substance. It is made up of 70% water, pancreatic and intestinal secretions, and desquamated epithelial cells from the intestinal tract and bile. It appears in the fetal intestine around 12 weeks of pregnancy and persists throughout the fetal colon during pregnancy. Almost all term babies pass their first stool by 48 hours after birth; meconium is typically passed in the first few hours. However, LBW babies typically have a delayed first stool passage(1–3).

A few theories have been put up to explain the Meconium’s journey through the foetus. In reaction to hypoxia, the foetus passes Meconium, which indicates foetal compromise. In utero passage of Meconium may represent normal gastrointestinal tract maturation under neural control. Meconium passage could follow vagal stimulation from common but transient umbilical cord entrapment and resultant increased peristalsis. It is masses of waste, desquamated cells from the intestine and skin, gastrointestinal mucin, lanugo fur, fatty material from the vernix caseosa, and intestinal secretions are all contained in Meconium-stained amniotic fluid(1,4,5).

Meconium is also a potential toxin if the fetus aspirates this particulate matter with gasping breath in utero or when it takes its first breaths following birth. Furthermore, inflammation of the lung, chorionic plate, and umbilical vessels is linked to intrauterine exposure to meconium, which increases perinatal morbidity(6–8).

Globally, the prevalence of intrapartum Meconium-stained amniotic fluid (MSAF) in term pregnancy ranges from 7% to 22 %(9). However, the incidence reaches up to 40% in a post-term pregnancy(7). One study in France found that 7.93% rate of term newborns had developed MSAF(10). According to studies conducted in various parts of India, 11.01 - 18 percent of laboring women have MSAF(5,11).

According to a report, MSAF affects 20.4 percent of the population in Nigeria(12) and 11.15% in Cameroon(13). In Ethiopia, the national pooled prevalence of MSAF shows 19.6%(14). A study done in Adama hospital medical college, the prevalence was 23.9%(15). In Saint Paul’s Hospital Millennium Medical College, Addis Ababa showed that, the Meconium was identified as grade I in (35%) grade-II in (33%), and grade-III (32%)(16). Other studies done in the district part of Ethiopia indicated that the magnitude of MSAF was 15.47%, 24.6% and 30.3% at Jimma University Specialized Teaching Hospital Felege Hiwot comprehensive specialized referral hospital and in Amhara Region North Shewa Zone hospitals, respectively(4,6,17).

It’s unclear exactly what causes amniotic fluid to be stained with meconium. Nonetheless, prior research indicated that obstetric variables such as prolonged labor, post-term pregnancy, low-birth weight babies, oligohydramnios, prolonged premature rupture of fetal membranes, intrauterine growth retardation, and hypertensive disorders of pregnancy, medical factors (Cholestasis of pregnancy and Anemia),and socio-demographic and behavioral risk factors (higher maternal age, maternal drug abuse especially tobacco, alcohol and cocaine use) are the major contributory factors for the passage of meconium into the amniotic fluid and Fetal factors include oligohydramnios, fetal growth restriction, and poor biophysical profile(3,12).

MSAF significantly increases the rate of maternal complications such as Meconium laden amniotic fluid embolism, intrapartum chorioamnionitis, puerperal endometritis, wound infection, increased risk of operative delivery, and its complication(6,7). Besides this, it increased rates of neonatal resuscitation, respiratory distress, lower APGAR score, neonatal nursery admissions, Meconium aspiration syndrome, neonatal sepsis, pulmonary disease and fetal-neonatal mortality(6,18–22).

If key risk factors are identified early, labour is properly monitored, and careful decisions are made regarding the timing and style of delivery, the prenatal morbidity and mortality associated with MSAF can be reduced(3).

The prevalence and predictors of Meconium-stained amniotic fluid are not known in the study area. Besides, all of the studies done in Ethiopia were conducted in a single referral hospital so the results might not be representative for other health care settings. In addition to this, other than the obstetric factors widely mentioned in literature, the current study also added variables like prolonged PROM, inter-pregnancy interval, and alcohol use that have been infrequently studied among Ethiopian studies of meconium stained amniotic fluid. It is critical to understand the local burden and factors associated with MSAF for better intrapartum care and neonatal surveillance. Therefore, this study aims to assess the prevalence of meconium-stained amniotic fluid and its associated factors among postnatal mothers in North Shewa Zone Public Hospitals, Oromia Ethiopia, 2023

## Methods and Materials

### Study design, setting and period

An institution based cross-sectional study design was conducted in North Shewa Zone, Oromia regional state, Ethiopia from February to March, 2023.

North Shewa Zone is one of the zones in the Oromia Region of Ethiopia. Its administrative city is Fitche town which is located 114km away from the capital city of Ethiopia, Addis Ababa.

In North Shewa Zone there are six public hospitals. Salale University Comprehensive Specialized Hospital, Kuyu General Hospital, Chancho Primary Hospital and Muka Turi Primary Hospital were hospitals in which the study was carried out.

### Source population

All women who gave birth in North Shewa Zone Hospitals were the source population.

### Study population

All women who gave birth at selected Hospitals in North Shewa Zone during data collection periods were considered as the study population.

### Eligibility criteria

All postnatal mothers who gave birth of singleton with >37weeks of gestational age and also who were permanent residents were included in the study and those mothers with Intra Uterine Fetal Death (IUFD), breech presentation, congenital anomaly, and who had in-complete chart were excluded from the study.

### Sample size determination

The sample size was calculated by a single population proportion formula n= (Z α/2)^2^ P (1- P)/ d^2^ based on the following assumptions: the prevalence of MSAF was 24.6 %(1), 95% confidence level of Z α/2= 1.96 and 0.05 margin of error. Thus, with 10% non-response rate gives the final sample which is **495**.

### Sampling procedure

From seven public hospitals four Hospitals were selected randomly using simple random sampling techniques. Salale university comprehensive specialized hospital, Kuyu General Hospital, Chancho Primary Hospital and Muka Turi Primary Hospital were selected. The sample size was proportionally allocated to each Hospitals based on their total number of one month of delivery coverage.

Dependent Variable:- Meconium-stained amniotic fluid

Independent variables:- Socio-demographic variables: age, residency, marital status, educational status, occupational status, and average monthly income.

Obstetrics and medical variable: Gestational age, ANC follow-up, Antepartum hemorrhage, Prolonged premature rupture of the membranes, oligohydraminos, Intrauterine growth restriction, Pregnancy-induced hypertension, Duration of labor > 15 hrs, Induction or augmentation, Previous Cesarean scar, Inter-pregnancy interval, Previous history of prenatal death, Chronic hypertension, and Maternal anemia.

Behavioral variables: alcohol consumption.

### Operational and term definition

Meconium-stained amniotic fluid: the presence of Meconium in the amniotic fluid changes the color of the liquor from clear to various shades of green, yellow or brownish color depending on the degree of Meconium-stained liquor(6).

Grade I MSAF: A small amount of Meconium is diluted in a plentiful amount of amniotic fluid. This gives the fluid only a slightly greenish or yellowish discoloration(6).

Grade II MSAF: is when there is a fair amount of amniotic fluid, but it is stained with Meconium. In this case, it will be ‘khaki green’ or brownish(6).

Grade III MSAF: Heavy staining, when there is reduced amniotic fluid and a large amount of Meconium, makes the staining quite thick, with a ‘pea soup’ consistency(6).

APGAR score: is determined by evaluating the new born baby on five simple criteria on a scale from zero to two, then summing up the five values thus obtained(6).

Gestational Age: Gestational age can be calculated by using first-trimester ultrasound, last normal menstruation period, and abdominal tap measurement(2).

Anemia during pregnancy: Those women who have a Hemoglobin level < 11.0 gm/ dl(23). Current alcohol use: defined as any use of alcohol in the past 30 days(24,25).

Prolonged premature rupture of membranes (PROM): the leakage of amniotic fluid that persists for more than 18 hours before the onset of labor(26).

### Data collection tools and procedures

Data were collected using a combination of interviews and chart review by four BSc midwives and two supervisors who were trained for this purpose. Structured interviewer-administered data collection questionnaires was adopted and modified from different kinds of literatures(1,4). Questionnaires that guided chart review and interview was structured into four logical sections (socio-demographic characteristics, obstetric related factors, medical history, and Behavioral related factors). Interview was conducted after informed written consent was taken from each participants. Data were collected by four BSc midwives and supervised by two Msc midwives.

### Data quality assurance

In order to maintain the quality of data, the questionnaire is developed in English and then it was translated to Afan Oromo and back to English. Before the actual data collection commences, the training was given to the data collectors and supervisor. Besides, the A Pretest was conducted on the 5% of the total sample sizes in the hospital found in North Shewa Zone, but not selected for study prior to the commencement of actual data collection. Data completeness was checked daily by the supervisor and principal investigator. The reliability of the tool was checked. Moreover, completeness and consistency of the collected data were reviewed and checked by supervisors and investigators.

### Data analysis procedures

The collected data were cleaned, coded and entered to the Epi Data version3.1, then exported to SPSS version 25. The frequency, proportion, means, and standard deviation was used to describe the data.

Descriptive data was presented by table, graphs, charts and means. Multiple logistic regression was used for variables with p value of < 0.25 in the bivariate analysis for not to miss the important variables. Odd Ratio and 95%CI was used to test strength of association at p-value of 0.05. Multicollinearity assumption was checked by the variance inflation factor (VIF) of < 1.6 which indicated that there is a less likely correlation between independent variables. The goodness of fit of the models was tested by using the Hosmer-Lemeshow test (p-value 0.381) and It was a good fit.

## Results

### Socio-demographic characteristics of the respondents

A total of 495 participants were responded to the questionnaires which account for 100% responses. The study participants’ mean age was 27.8 (SD ± 6.034 years), with a minimum and maximum age of 18 and 44 years, respectively. Regarding place of residence 294 (59.4%) participants were urban residents. (Table**2**).

### Obstetric and medical-related characteristics

Near to half 223 (45.1%) of the participants were multiparous. The majority of the participants 311 (62.8%) gestational age were between 37 weeks -42 weeks and the mean gestational age was 40.56 weeks and with an SD of ± 1.398 weeks. (Table 3)

**Table 1.**
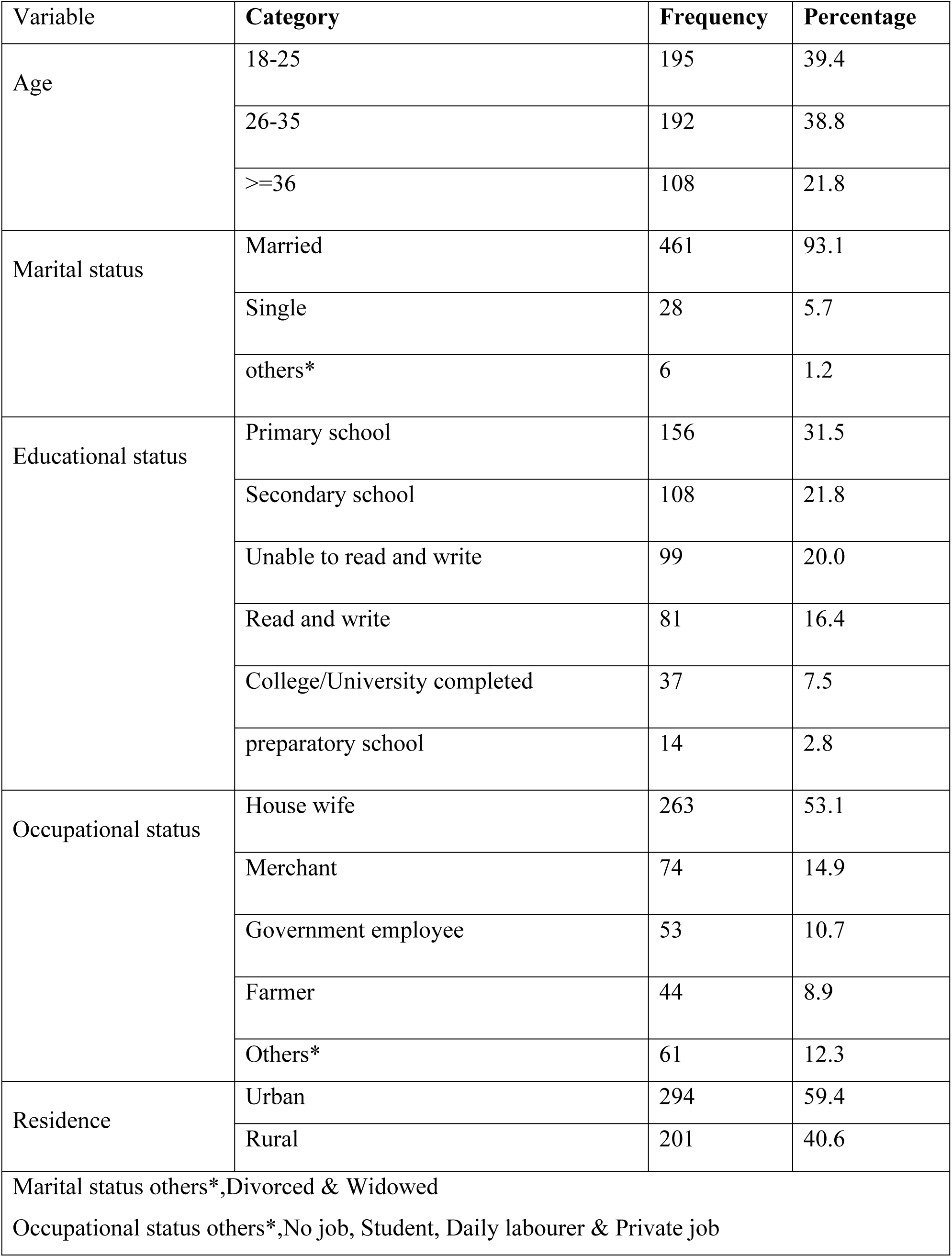
Socio-demographic characteristics of prevalence of Meconium-stained amniotic fluid and associated factors among postnatal mothers in North Shewa Zone Hospitals, 2023 (n=495)

**Table 2.** Obstetric and medical-related characteristics of postnatal index mothers North Shewa Zone Public Hospitals, Ethiopia 2022 (n=495)

| Variable | Category | Frequency | Percentage |
|---|---|---|---|
| Parity | Primiparous | 189 | 38.2 |
|  | Multiparous | 223 | 45.1 |
|  | Grand-multipra | 83 | 16.8 |
| Gestational age in weeks | Term(37 weeks-42 weeks) | 311 | 62.8 |
|  | post term(≥42 weeks) | 184 | 37.2 |
| ANC follow up | Yes | 462 | 93.3 |
|  | No | 33 | 6.7 |
| Number of ANC follow up(N=462) | one contact | 15 | 3.3 |
|  | 2-3 contacts | 99 | 21.4 |
|  | >=4 contacts | 348 | 75.3 |
| Maternal Rh status | Positive | 482 | 97.4 |
|  | Negative | 13 | 2.6 |
| History of Antepartum hemorrhage | Yes | 267 | 53.9 |
|  | No | 228 | 46.1 |
| Prolonged Premature rupture of the Membranes | Yes | 205 | 41.4 |
|  | No | 290 | 58.6 |
| Intrauterine growth restriction | Yes | 177 | 35.8 |
|  | No | 318 | 64.2 |
| Pregnancy-induced Hypertension | Yes | 105 | 21.2 |
|  | No | 390 | 78.8 |
| Duration of labor > 15 hrs. | Yes | 184 | 37.2 |
|  | No | 311 | 62.8 |
| Obstructed labor | Yes | 111 | 22.4 |
|  | No | 384 | 77.6 |
| Induction or augmentation | Yes | 199 | 40.2 |
|  | No | 296 | 59.8 |
| Current pregnancy Mode of delivery | Spontaneous vaginal delivery | 374 | 75.6 |
|  | Cesarean section | 77 | 15.6 |
|  | Instrumental delivery | 44 | 8.9 |
| Previous Cesarean scar(N=306) | Yes | 23 | 7.5 |
|  | No | 283 | 92.5 |
| Inter pregnancy interval (N=285) | <2years | 84 | 29.5 |
|  | >5years | 201 | 70.5 |
| Previous history of prenatal death(N=306) | Yes | 47 | 15.4 |
|  | No | 259 | 84.6 |
| Oligo-hydraminous | Yes | 187 | 37.8 |
|  | No | 308 | 62.2 |
| Maternal anemia | Yes | 68 | 13.7 |
|  | No | 427 | 86.3 |
| Chronic hypertension | Yes | 33 | 6.7 |
|  | No | 462 | 93.3 |

**Table 3.** Prevalence of Meconium-stained amniotic fluid and associated factors among postnatal mothers in North Shewa Zone Hospitals, Ethiopia 2023 (n=495)

| **Variable** | **Category** | **Frequency** | **Percentage** |
|---|---|---|---|
| **Nature of liquor** | Clear | 347 | 70.1 |
|  | Meconium | 148 | 29.9 |
| Time in which meconium-stained amniotic fluid stained noticed(N=148) | immediately at membrane rupture | 77 | 52.1 |
|  | Initial clear liquor followed by MSAF | 71 | 47.9 |

### Behavioral characteristics

From the total of participants, 172 (34.7%) were alcohol users for the last 30 days.

### Prevalence of meconium-stained amniotic fluid

The prevalence of Meconium-stained amniotic fluid was found 29.9% (95%CI; 25.9-34.1). From that of meconium stained liquor, 77(52.1%) were noted immediately at membrane rupture. (Table 4)

**Table 4.**
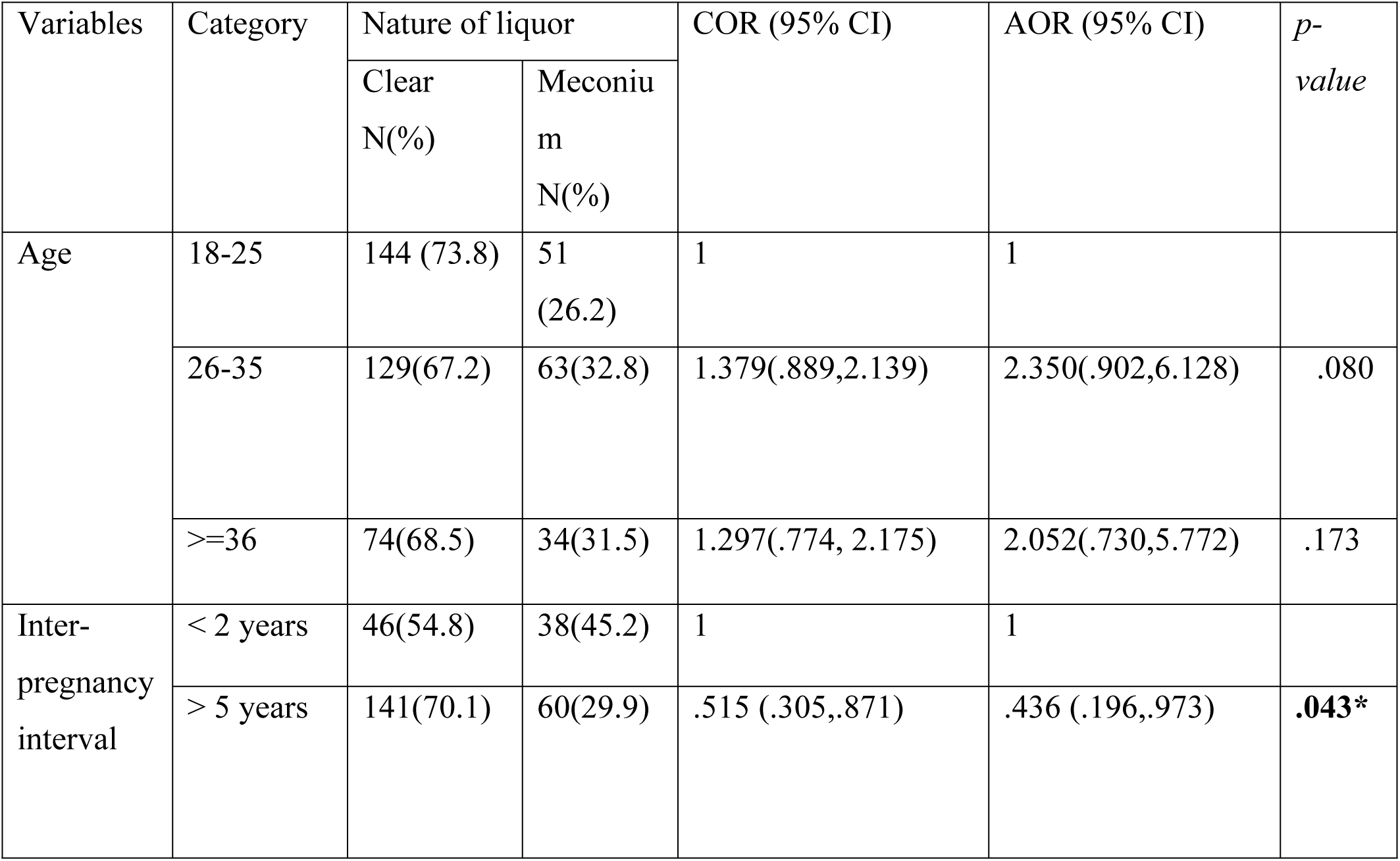
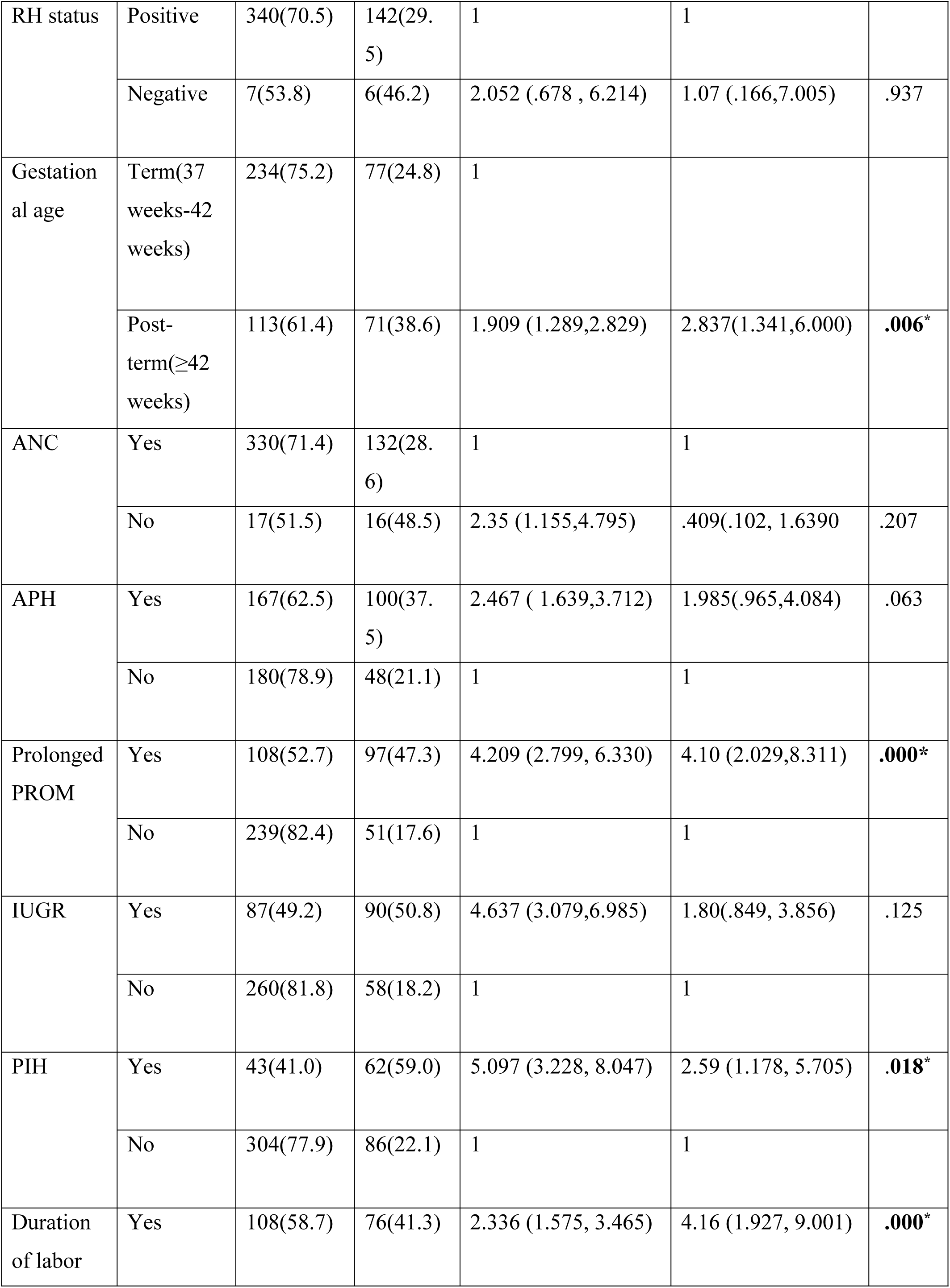
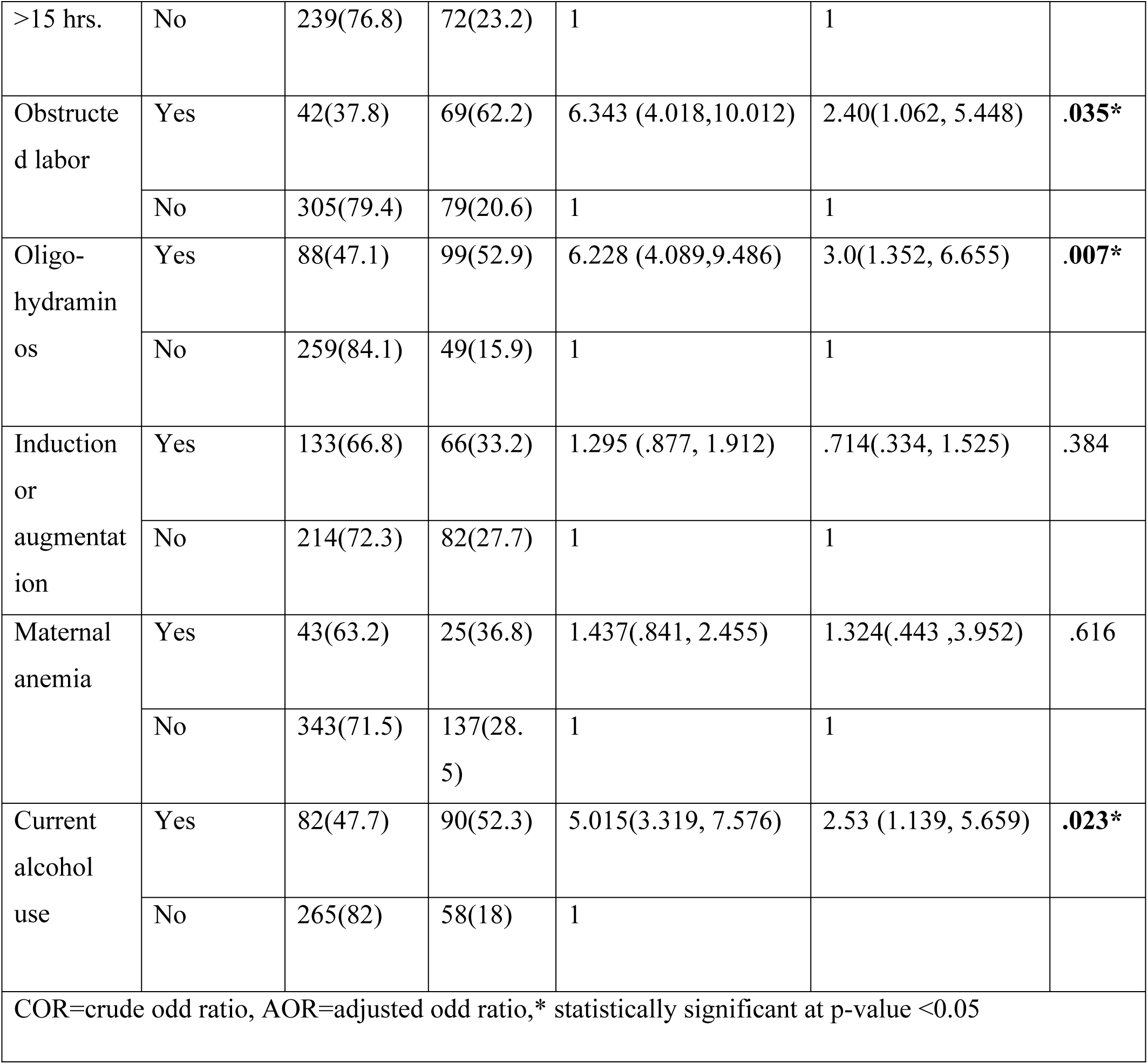
Binary and Multiple logistic regression results for factors associated of factors of Meconium-stained amniotic fluid among postnatal mothers in North Shewa Zone hospitals, Ethiopia 2023 (n=495)

| Variables | Category | Nature of liquor |  | COR (95% CI) | AOR (95% CI) | *p-value* |
|---|---|---|---|---|---|---|
|  |  | Clear<br>N(%) | Meconium<br>N(%) |  |  |  |
| Age | 18-25 | 144 (73.8) | 51 (26.2) | 1 | 1 |  |
|  | 26-35 | 129(67.2) | 63(32.8) | 1.379(.889,2.139) | 2.350(.902,6.128) | .080 |
|  | >=36 | 74(68.5) | 34(31.5) | 1.297(.774, 2.175) | 2.052(.730,5.772) | .173 |
| Inter-pregnancy interval | < 2 years | 46(54.8) | 38(45.2) | 1 | 1 |  |
|  | > 5 years | 141(70.1) | 60(29.9) | .515 (.305,.871) | .436 (.196,.973) | **.043\*** |
| RH status | Positive | 340(70.5) | 142(29.5) | 1 | 1 |  |
|  | Negative | 7(53.8) | 6(46.2) | 2.052 (.678 , 6.214) | 1.07 (.166,7.005) | .937 |
| Gestational age | Term(37 weeks-42 weeks) | 234(75.2) | 77(24.8) | 1 |  |  |
|  | Post-term(≥42 weeks) | 113(61.4) | 71(38.6) | 1.909 (1.289,2.829) | 2.837(1.341,6.000) | **.006**<sup>\*</sup> |
| ANC | Yes | 330(71.4) | 132(28.6) | 1 | 1 |  |
|  | No | 17(51.5) | 16(48.5) | 2.35 (1.155,4.795) | .409(.102, 1.6390 | .207 |
| APH | Yes | 167(62.5) | 100(37.5) | 2.467 ( 1.639,3.712) | 1.985(.965,4.084) | .063 |
|  | No | 180(78.9) | 48(21.1) | 1 | 1 |  |
| Prolonged PROM | Yes | 108(52.7) | 97(47.3) | 4.209 (2.799, 6.330) | 4.10 (2.029,8.311) | **.000\*** |
|  | No | 239(82.4) | 51(17.6) | 1 | 1 |  |
| IUGR | Yes | 87(49.2) | 90(50.8) | 4.637 (3.079,6.985) | 1.80(.849, 3.856) | .125 |
|  | No | 260(81.8) | 58(18.2) | 1 | 1 |  |
| PIH | Yes | 43(41.0) | 62(59.0) | 5.097 (3.228, 8.047) | 2.59 (1.178, 5.705) | **.018**<sup>\*</sup> |
|  | No | 304(77.9) | 86(22.1) | 1 | 1 |  |
| Duration of labor >15 hrs. | Yes | 108(58.7) | 76(41.3) | 2.336 (1.575, 3.465) | 4.16 (1.927, 9.001) | **.000**<sup>\*</sup> |
|  | No | 239(76.8) | 72(23.2) | 1 | 1 |  |
| Obstructed labor | Yes | 42(37.8) | 69(62.2) | 6.343 (4.018,10.012) | 2.40(1.062, 5.448) | **.035\*** |
|  | No | 305(79.4) | 79(20.6) | 1 | 1 |  |
| Oligo-hydraminos | Yes | 88(47.1) | 99(52.9) | 6.228 (4.089,9.486) | 3.0(1.352, 6.655) | **.007\*** |
|  | No | 259(84.1) | 49(15.9) | 1 | 1 |  |
| Induction or augmentation | Yes | 133(66.8) | 66(33.2) | 1.295 (.877, 1.912) | .714(.334, 1.525) | .384 |
|  | No | 214(72.3) | 82(27.7) | 1 | 1 |  |
| Maternal anemia | Yes | 43(63.2) | 25(36.8) | 1.437(.841, 2.455) | 1.324(.443 ,3.952) | .616 |
|  | No | 343(71.5) | 137(28.5) | 1 | 1 |  |
| Current alcohol use | Yes | 82(47.7) | 90(52.3) | 5.015(3.319, 7.576) | 2.53 (1.139, 5.659) | **.023\*** |
|  | No | 265(82) | 58(18) | 1 |  |  |

COR=crude odd ratio, AOR=adjusted odd ratio,\* statistically significant at p-value <0.05

### Grade of meconium stained amniotic fluid

Regarding the grade of meconium stained amniotic fluid status, the majority, and 54(36.4%) was grade II. (**Fig 1**)

**Figure 1.**
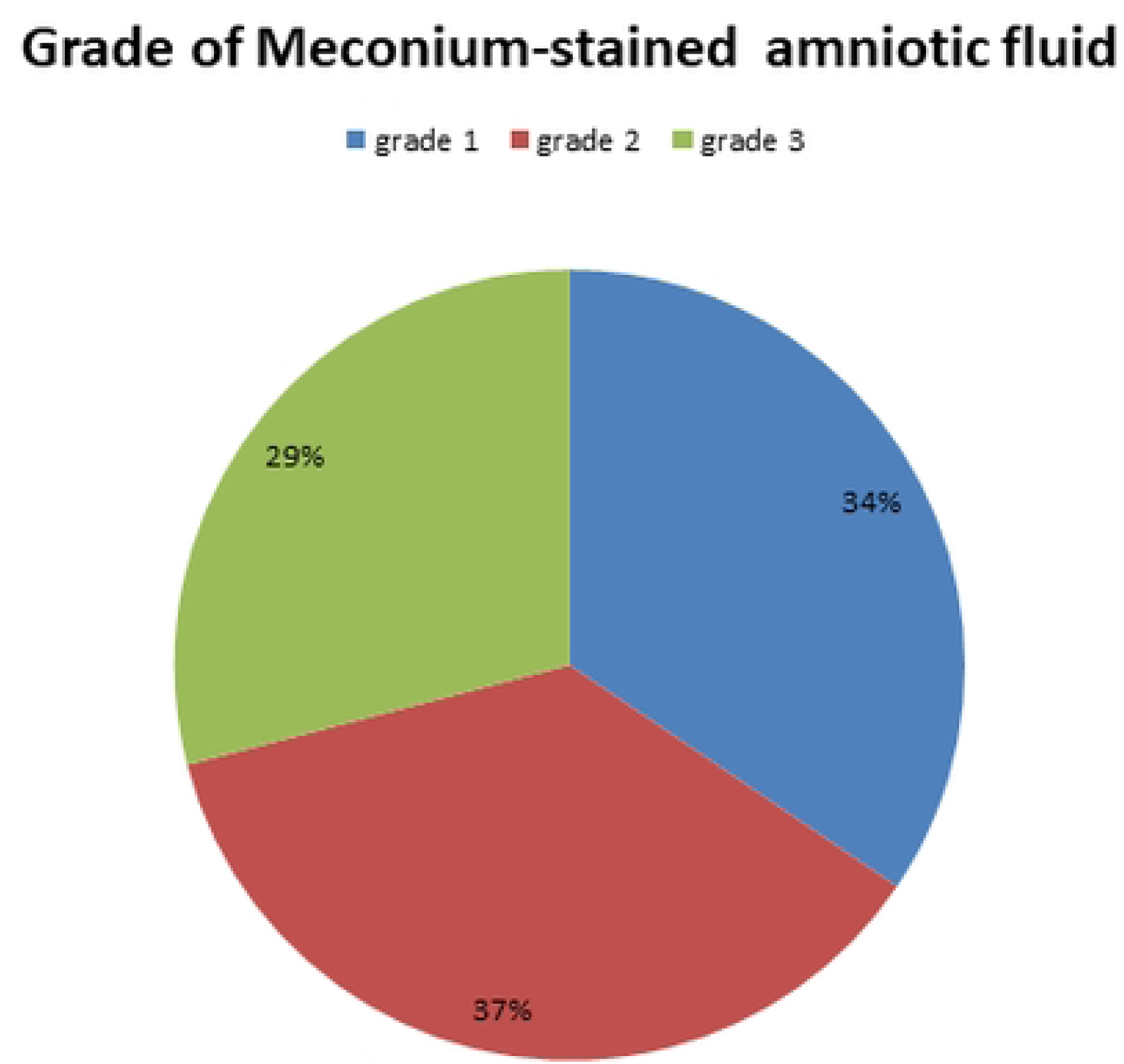
Grade of Meconium-stained amniotic fluid among postnatal mothers in North Shewa Zone

### Factors associated with meconium-stained amniotic fluid

Inter-pregnancy interval, Gestational age, Prolonged PROM, PIH, Duration of labor >15 hrs, Obstructed labor, Oligohydraminos and Current alcohol use were significantly associated with Meconium stained amniotic fluid at a p-value ≤ 0.05.

As Compared to women who had an inter-pregnancy interval of less than 2 years, those with an interval greater than 5 years had 56.4% lower odds of MSAF [AOR = .436, 95%CI =.196,.973]. Mothers with a post-term Gestational age had about 2.8 times higher odds of developing MSAF than those term pregnancies. [AOR = 2.837, 95%CI = 1.341, 6.000]. Mothers who had been diagnosed with Prolonged PROM were 4.1 times more likely to develop MSAF during labor as compared to those who don’t develop prolonged PROM [AOR = 4.10, 95%CI = 2.029,8.311]. Those women with a duration of labor > 15 hrs had about 4.1 times higher odds of developing MSAF than those having less than 15 hrs [AOR = 4.16, 95%Cl =1.927, 9.001]. Finally, The chance of developing MSAF in mothers whose diagnosed with maternal oligo-hydraminos was 3 times higher than in those having normal amniotic fluid volume [AOR = 3.0, 95%CI = 1.352, 6.655]. Mothers with obstructed labor had about 2.4 times higher odds of developing MSAF. [AOR = 2.40, 95%CI = 1.062, 5.448]. Those women who had been current alcohol users had about 2.53 times higher odds of developing MSAF [AOR = 2.53, 95%Cl =1.139, 5.659]. (Table 5)

## Discussion

This study found that the proportion of Meconium-stained amniotic fluid was found to be 29.9% (95%CI; 25.9-34.1). The prevalence observed in this study was higher than findings reported from several Ethiopian and international studies. local reports which were done in Ethiopian regions shows the proportion of Meconium-stained amniotic fluid were 23.9%, 24.6%, 15.4% and 17.8% respectively(1,4,6,15). The finding was higher than the studies from African countries i.e Nigeria 20%(12) and Cameroon 11.5(13). The incidence of Meconium-stained liquor from different studies done in India shows 18%(11) and 11%(5) which made our finding higher. It is also higher than studies from France 8.2% (27) and Brazil 11.9% (28).

This disparity could be caused by a number of socioeconomic factors, variations in study populations, obstetric characteristics, including the accessibility & utilization of healthcare services, referral patterns and the quality of intrapartum monitoring. Some of the studies were studied at single institutions and some used a method of retrospective data collection from records. Moreover, the area in which we conducted this study was affected by security instability, which had a significant impact on all aspects of the residents’ lives, including social and economic ones, mainly the utilization of healthcare services and the timely referral patterns. The relatively high prevalence observed in this study may indicate the need for strengthened fetal surveillance during labor and timely management of obstetric complications.

This study prevalence was comparable with a study conducted in North Shewa Zone, Amhara Region, which reported a prevalence of 30.3%(17). The similarity in findings may be related to similarities in the study population, geographical setting, and maternal and obstetric characteristics of the participants.

The multivariate logistic regression result revealed that, the chance of developing MSAF in mothers whose diagnosed with maternal Oligo-hydraminos was 2.5 times higher than in those having normal amniotic fluid volume. Our result finding is supported with different studies conducted in Ethiopia (1,4) and India(8,18). A possible explanation for the reason could be, reduced amniotic fluid volume may reflect placental insufficiency and chronic fetal compromise, which can increase fetal stress and stimulate meconium passage before delivery(29).

Duration of labor greater than 15 hours had about 4.4 times higher odds of developing MSAF than those having less than 15 hrs. this find had been supported by different studies(1,4,17) and India (13,18). Prolonged labor may expose the fetus to sustained stress and intermittent hypoxia, resulting in increased intestinal peristalsis and relaxation of the anal sphincter, allowing meconium to slip through(30). This finding highlights the importance of close labor monitoring and early identification of abnormal labor progress.

Similarly, prolonged premature rupture of membranes was significantly associated with MSAF. Mothers with prolonged PROM had substantially higher odds to develop MSAF than those without prolonged PROM during labor. One possible explanation is that prolonged rupture of membranes increases the risk of intrauterine infection and fetal compromise, both of which may contribute to meconium passage before birth. This could ultimately have an impact on the health of the foetus(31,32).

Inter-pregnancy interval was significantly associated with MSAF. Mothers with an interval greater than 5 years had 56.4% lower odds of MSAF than who had an interval of less than 2 years.

This implies that there has been less time for the mother’s body to recover its micronutrients and repair the endometrium tissue. It is mostly used to study along with preterm labor and low birth weight because short interval period indirectly leads to MSAF through placental problems and growth restrictions(33). Although the underlying mechanism is not clear, further research is required to clarify this relationship.

post-term gestational age was also independently associated with MSAF. Mothers with a post-term Gestational age had about 2.8 times higher odds of developing MSAF. The increased risk of MSAF in increasing gestational age may be due to age-related changes in the placenta that lead to decreased placental reserve and may cause fetal stress leading to passage of meconium. In post-term pregnancy (> 42 weeks), the gastrointestinal tract of fetus is fully developed, and physiological level of motilin increases, thus leading to passage of meconium in the absence of fetal distress(34,35).

Hypertension during pregnancy was linked to a 2.6 times increased risk of MSAF. PIH, including preeclampsia cause injury to the placental blood vessels and resulting in decreased oxygen delivery to the fetus. The fetus responds physiologically through the vagal nerve by contracting the intestinal tract and relaxing the anal sphincter. In general the pathophysiology of hypertensive disorders can cause impaired uteroplacental and fetal oxygenation, increasing the risk of meconium passage(36).

This finding is consistent with the Felege Hiwot study, which reported an independent association between preeclampsia and MSAF. This was congruent with a study conducted at St Paul’s Hospital Millennium Medical College, Addis Ababa, Ethiopia(16) and Felege Hiwot comprehensive specialized referral hospital, North West Ethiopia(4).

Mothers with obstructed labor had about 2.4 times higher odds of developing MSAF. In cases where a woman experiences obstructed labor, the delay becomes increasingly harmful to the baby due to its mechanical nature. Obstetric difficulties during labor may affect the utero-placental perfusion. Prolonged uterine contractions will lead to umbilical cord and placenta compression and ultimately hypoxemia and acidosis and cause fetal stress, potentially resulting in passage of meconium(37).

This finding is consistent with the studies done from different parts of Ethiopia(1,4,17,37), which also identified obstructed labor as an independent predictor of MSAF.

Finally, current alcohol use was associated with about 2.53 times higher odds of developing MSAF. Maternal substance use, including alcohol, acts as a lifestyle risk factor. Alcohol affects oxygen delivery through the placenta. Even if it increases the baseline statistics for MSAF, it generally ranks lower than acute condition like PIH(24).

These research findings have major clinical implications. Because of the relatively high prevalence of MSAF, there should be intensified intrapartum monitoring and detection of mothers at risk like those with prolonged labor, oligohydramnios, and prolonged PROM. Proper management of labor, immediate obstetric intervention, and neonatal resuscitation readiness may lessen complications that may arise from MSAF.

## Conclusions

The prevalence of MSAF was relatively high in the study area. Prolonged premature rupture of membranes, duration of labor greater than 15 hours, current alcohol use, obstructed labor, PIH, post-term gestational age, Inter-pregnancy interval and oligohydramnios were independently associated with MSAF. These findings highlight the importance of identifying maternal and obstetric risk factors and strengthening appropriate intrapartum assessment and management. Women with hypertension during pregnancy, abnormal labor, and other recognized risk factors require more consideration as their need for obstetrical care and neonatal resuscitation is higher.

It is more beneficial for the government and the health bureau in the area to organize proper health education for the mothers in pre-conception care and antenatal care as far as early detection and prevention of diseases are concerned. In addition to this, it is necessary to promote continuous professional development for maternity care providers on the identification and management of obstetric complications associated with MSAF.

Healthcare workers need to strengthen antenatal and intrapartum monitoring in order to allow early detection, careful monitoring, and management of women who have risks that predispose to the development of MSAF. Early and prolonged detection by use of a partograph and latent phase chart, as well as prompt intervention, is encouraged. Screening on alcohol consumption and counseling needs to be incorporated in antenatal visits, although the link established in this study still needs validation.

It is recommended that additional research studies be carried out to establish the cause-and-effect relationship as well as assess the neonatal outcomes linked to the MSAF. Future Investigate the connection between alcohol intake by mothers and the MSAF using more detailed exposure measurements and appropriate control of potential confounders.

## Limitations of the study

The cross-sectional design restricts the possibilities of establishing temporal connections and causality between the factors and MSAF. Thus, the associations that have been found should be treated as statistical connections rather than causations.

Alcohol intake was evaluated using self-reports, and the results may be influenced by recall bias and social desirability bias, potentially leading to underreporting of alcohol use.

## Strength

The possibility that more than one hospital was used may increase the representativeness of the findings in comparison with those of other studies conducted in a single hospital. In addition to the obstetric factors usually researched, the study included the analysis of such risk factors for MSAF as alcohol use status and inter-pregnancy interval.

## Ethics statement

This project was conducted after getting approval by the Salale University College of medicine and health science Institutional Review Board (IRB). Data were collected after getting official permission from North Shewa Zone health bureau and each selected Hospitals administration office. All study participants were informed about the purpose, the risk, the benefits of the study, their right to refuse participation and written voluntary consent was obtained from all study participants before the interview. The respondents were also informed that the information obtained was treated with extreme confidentiality.

## Abbreviations

ANC: Ante Natal Care
CS: Cesarean Section
DM: Diabetic Mellitus
GA: Gestational Age
IUFD: Intra-Uterine Fetal Demise
IUGR: Intra Uterine Growth Restriction
MAS: Meconium Aspiration Syndrome
MSAF: Meconium Stained Amniotic Fluid
MSL: Meconium Stained Liquor
NICU: Neonatal Intensive Care Unit
OPD: Out Patient Diagnosis
OR: Odds Ratio
PIH: Pregnancy Induce Hypertension
PROM: Premature Rupture of Membrane
WHO: World Health Organization

## Notes

### Competing Interest Statement

The authors have declared no competing interest.

